# IDR-induced CAR condensation improves the cytotoxicity of CAR-Ts against low-antigen cancers

**DOI:** 10.1101/2023.10.02.560460

**Authors:** Xinyan Zhang, Qian Xiao, Longhui Zeng, Fawzaan Hashmi, Xiaolei Su

## Abstract

Chimeric antigen receptor (CAR)-T cell-based therapies demonstrate remarkable efficacy for the treatment of otherwise intractable cancers, particularly B-cell malignancies. However, existing FDA-approved CAR-Ts are limited by low antigen sensitivity, rendering their insufficient targeting to low antigen-expressing cancers. To improve the antigen sensitivity of CAR-Ts, we engineered CARs targeting CD19, CD22, and HER2 by including intrinsically disordered regions (IDRs) that promote signaling condensation. The “IDR CARs” triggered enhanced membrane-proximal signaling in the CAR-T synapse, which led to an increased release of cytotoxic factors, a higher killing activity towards low antigen-expressing cancer cells in vitro, and an improved anti-tumor efficacy in vivo. No elevated tonic signaling was observed in IDR CAR-Ts. Together, we demonstrated IDRs as a new tool set to enhance CAR-T cytotoxicity and to broaden CAR-T’s application to low antigen-expressing cancers.

## Introduction

T cells engineered with the chimeric antigen receptor (CAR) emerge as a versatile and effective toolset for cancer immunotherapy. CAR-Ts have been implemented in combating a variety of cancers and have achieved unprecedented success in treating blood malignancy (Ghobadi, 2018; Lee et al., 2015; Maldini et al., 2018; Porter et al., 2011). By 2023, there are six FDA-approved CAR-T therapies for certain types of leukemia, lymphoma, and myeloma. These achievements encouraged further development of CAR-Ts targeting an expanding pool of cancerous, infectious, autoimmune and fibrotic diseases (Aghajanian et al., 2019; Amor et al., 2020; Elinav et al., 2008; Ellebrecht et al., 2016; Hale et al., 2017; Leibman et al., 2017).

Meanwhile, CAR-T therapy is still limited by a few major obstacles, one of which is limited signaling efficiency. Though derived from the T cell receptor (TCR), CAR displays a much lower antigen sensitivity than TCR: a few hundred or more antigen molecules are required to activate a CAR-T cell (Gudipati et al., 2020; Majzner et al., 2020; Watanabe et al., 2015) whereas a single peptide-loaded MHC molecule is sufficient to trigger the activation of a normal T cell (Huang et al., 2013; Irvine et al., 2002). This low antigen sensitivity not only narrows CAR-T’s target to high antigen-expressing cancers, but also brings a challenge in maintaining sustained CAR-T activity against tumors; CAR T-treated patients experience an up to 60% of relapse, which is mostly caused by antigen loss (Majzner and Mackall, 2018; Maude et al., 2018; Park et al., 2018). Therefore, there is a critical need to develop CAR-Ts that can respond to low antigen-expressing cancer cells.

Mechanistically, the cause for low antigen sensitivity of CAR remains incompletely understood. The binding of antigen to CAR is much tighter (with a Kd of nM to pM) than the binding of pMHC to TCR (with a Kd of μM) and therefore, the inability of CAR-Ts to be activated by low antigen-expressing cells more likely results from the signal processing mechanism downstream CAR rather than the binding of CAR to antigen. Previous studies suggested that CARs insufficiently induce membrane-proximal signaling including ZAP70 activation and LAT phosphorylation, and LAT is weakly engaged in the CAR pathway (Dong et al., 2020; Gudipati et al., 2020; Salter et al., 2021). LAT enhances TCR signal transduction by promoting the macromolecular complex assembly and condensation of TCR signaling molecules (Houtman et al., 2006; Huang et al., 2019; Su et al., 2016; Zeng et al., 2021). These suggested an opportunity to engage condensation to improve CAR-T signaling, especially on its response to low antigen-expressing cancers.

Intrinsically disordered regions (IDRs) received increasing attention because of their ability to form biomolecular condensates. IDRs do not typically fold into a well-defined 3D structure. Instead, they form condensates via weak inter and intra molecular interactions (Borcherds et al., 2021; Pappu et al., 2023). These condensates display unique biochemical activities through enriching and organizing effector molecules to promote cell signaling (Case et al., 2019; Xiao et al., 2022a). Here we decided to induce CAR condensation through constructing CAR-IDR fusion proteins. We identified a few IDRs, including those from FUS, EWS, and TAF15, that promote the membrane-proximal signaling, cytotoxic factor production, and killing of CAR-Ts against low antigen-expressing cancers (CD19, CD22, and HER2). Moreover, no elevated tonic signaling was observed in IDR CAR-Ts, suggesting a difference in signaling outcomes between IDR-induced condensation and previously reported CAR aggregation (Chen et al., 2023; Long et al., 2015). Together, these results demonstrated that IDRs, though not originally linked to T cell signaling, can serve as a new modular motif to improve the anti-tumor effect of CAR-Ts.

## Results

### Construction of IDR CAR-Ts

IDRs contain diverse sequence and structure features. To determine which IDR promotes the condensation of CAR, we selected candidates from 6 well-characterized IDRs that were previously shown to induce condensation in a cellular environment. These include IDRs from FUS, EWS, TAF15, Nup98, TDP43, and a synthetic IDR (synIDR) (Chong et al., 2018; Dzuricky et al., 2020; Kato et al., 2012; Protter et al., 2018; Wang et al., 2018; Xu et al., 2021; You et al., 2020). We chose CD19 CAR, which is commonly used in research and clinical practice, as an initial model. This CAR is composed of a single chain variable fragment (scFv, FMC63) that targets CD19, a stalk and transmembrane domain from CD8α, and cytosolic signaling domains from 41BB, CD28, and CD3ζ. The IDR was fused to the C-terminus of CD3ζ. A superfolder GFP tag, which promotes the folding of fused client proteins (Pedelacq et al., 2006), was further attached on the C-terminus of IDR for visualization of CAR condensation (Fig. 1A). The superfolder GFP tag enables live cell imaging, which avoids potential fixation-induced artifacts in characterizing IDR condensation (Irgen-Gioro et al., 2022). The DNA fragment encoding the control or IDR CAR was packaged into lentivirus and introduced into primary CD3^+^ T cells purified from the human peripheral mononuclear cells (PBMCs). Flow cytometry revealed the total cellular expression (by GFP) versus cell surface localization (by FMC63) of individual CARs (Fig. 1B), demonstrating that fusion with IDR did not affect the trafficking of CAR to the cell surface. To visualize the condensation of CAR on the cell surface, we stained live CAR-T cells with a plasma membrane dye CellMask deep red and performed total internal reflection fluorescence (TIRF) microscopy which effectively reduces the cytosolic background of fluorescence. We found that the CAR fused with FUS, EWS, or TAF15 displayed an enhanced condensation as compared to the control CAR (Fig. 1C, 1D). Therefore, we focused on these three CARs in the following functional assays.

**Figure 1.**
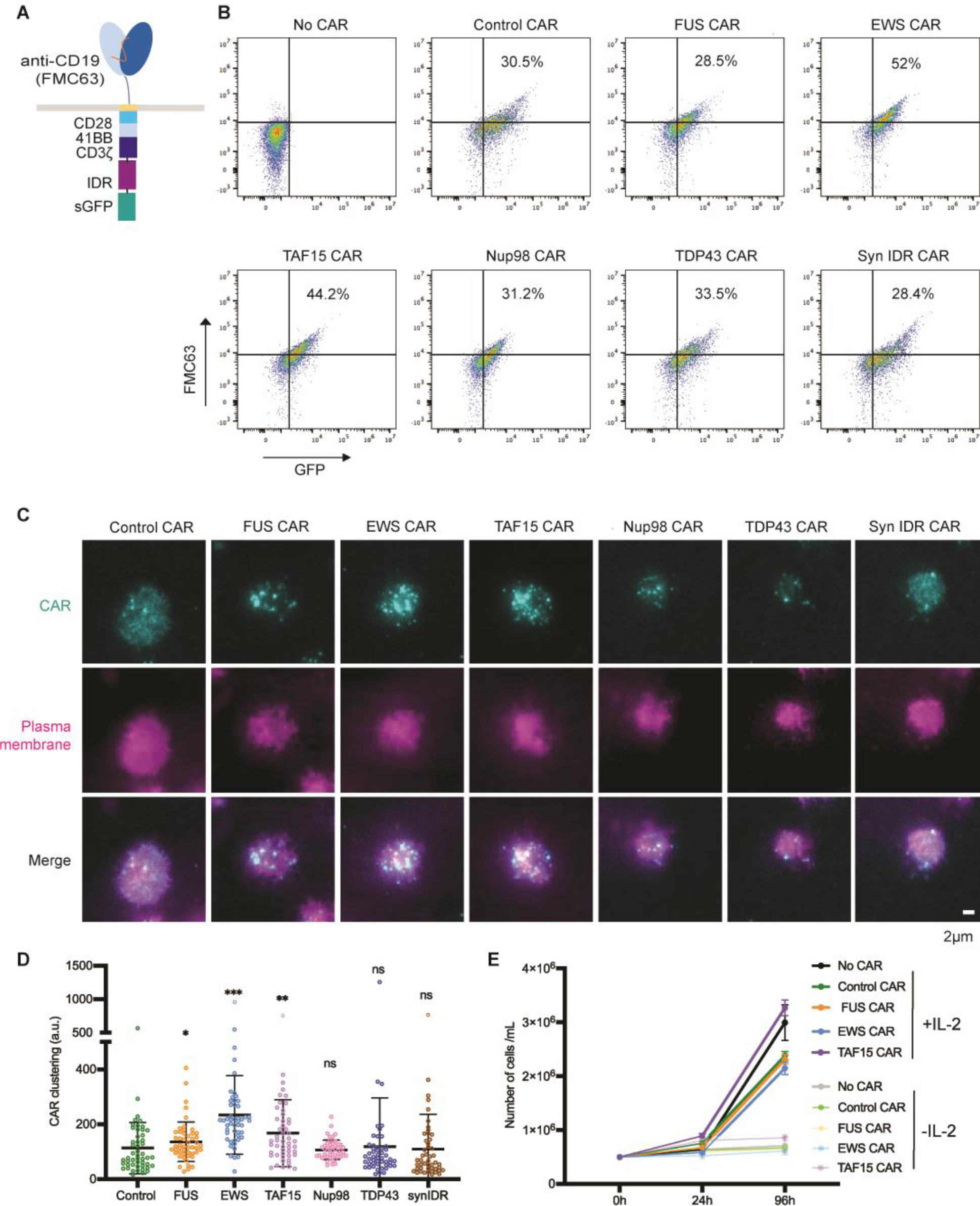
IDR promoted CAR condensation on the T cell surface. A) Schematics of the CD19 CAR used in this study. It contained an scFv targeting CD19 (FMC63), a CD8 hinge and transmembrane domain, an intracellular signaling domain composed of CD28, 41BB, and CD3ζ, an intrinsically disordered region (IDR), and superfolder GFP. B) Expression of IDR CARs. Human pan primary T cells were transduced with lentivirus encoding the control or IDR CAR. The expression level of CAR on the T cell surface was measured via flow cytometry using an anti-FMC63 antibody. A GFP tag was fused on the C-terminus of each CAR to monitor the total expression level of CAR. C) Condensation of CAR on the plasma membrane. TIRF microscopy revealed condensation of the control or IDR CAR on the T cell membrane. GFP was tagged on the C-terminus of CAR for visualizing CAR. The plasma membrane was labeled by a CellMask Deep Red dye. D) Quantification of CAR clustering by normalized variance. N=48-50 cells. Shown are mean ± std. All comparisons in this and following figures were made between individual IDR CARs and the control CAR (n.s. p>=0.05, *\* p<0.05, ** p<0.01, *** p<0.001*). E) CAR-T cell proliferation with or without IL-2. Quantification of the absolute cell number in the culture with or without IL-2. N=3 independent experiments. Shown are mean ± std.

Previous work showed that aggregation of CARs targeting GD2 or CSPG4 induces tonic signaling, which triggers CAR-T activation in the absence of antigens (Chen et al., 2023; Long et al., 2015). Therefore, we assessed if IDR-induced CAR condensation causes tonic signaling. We found that the IDR CAR-Ts did not proliferate in the absence IL-2 (Fig. 1E). The expression of CD69, a T cell activation marker, was similar between the control and IDR CAR-Ts in the absence of antigen (Fig. S1A). Moreover, IDR CAR-Ts did not secret detectable TNFα in the absence of antigen (Fig. S1B). Together, assessed from cell proliferation, activation marker, and cytokine production, IDR-induced CAR condensation did not trigger tonic signaling.

### IDR enhanced the cytotoxicity of CD19 CAR-T

To determine how IDRs affect the cytotoxicity of CAR-Ts against cancer cells, we co-cultured the control or IDR CAR-Ts (Fig. 2A) with variants of Nalm6, a B cell leukemia line, that expresses either high or low CD19 (Fig. 2B). The Nalm6 cells express a luciferase reporter which enables the quantification of cytotoxicity by the luciferase assay. The FUS and EWS, but not TAF15 CAR, displayed a substantial higher cytotoxicity towards both CD19-high and low Nalm6 cells (Fig. 2C and 2D). This result was recapitulated using another B cell line Raji as the target (Fig. S2A - S2C). The higher cytotoxicity of FUS and EWS CAR could be explained by their higher secretion of cytotoxic factors including Granzyme A, Granzyme B, Perforin, FASL, and IFNꝨ, when CAR-Ts were engaged with CD19-low Nalm6 cells (Fig. 2E-2I and S2D). Together, these data showed that the FUS and EWS IDRs enhanced the cytotoxicity of CAR-Ts against CD19-low cells, which was accompanied with a higher secretion of cytotoxic factors.

**Figure 2.**
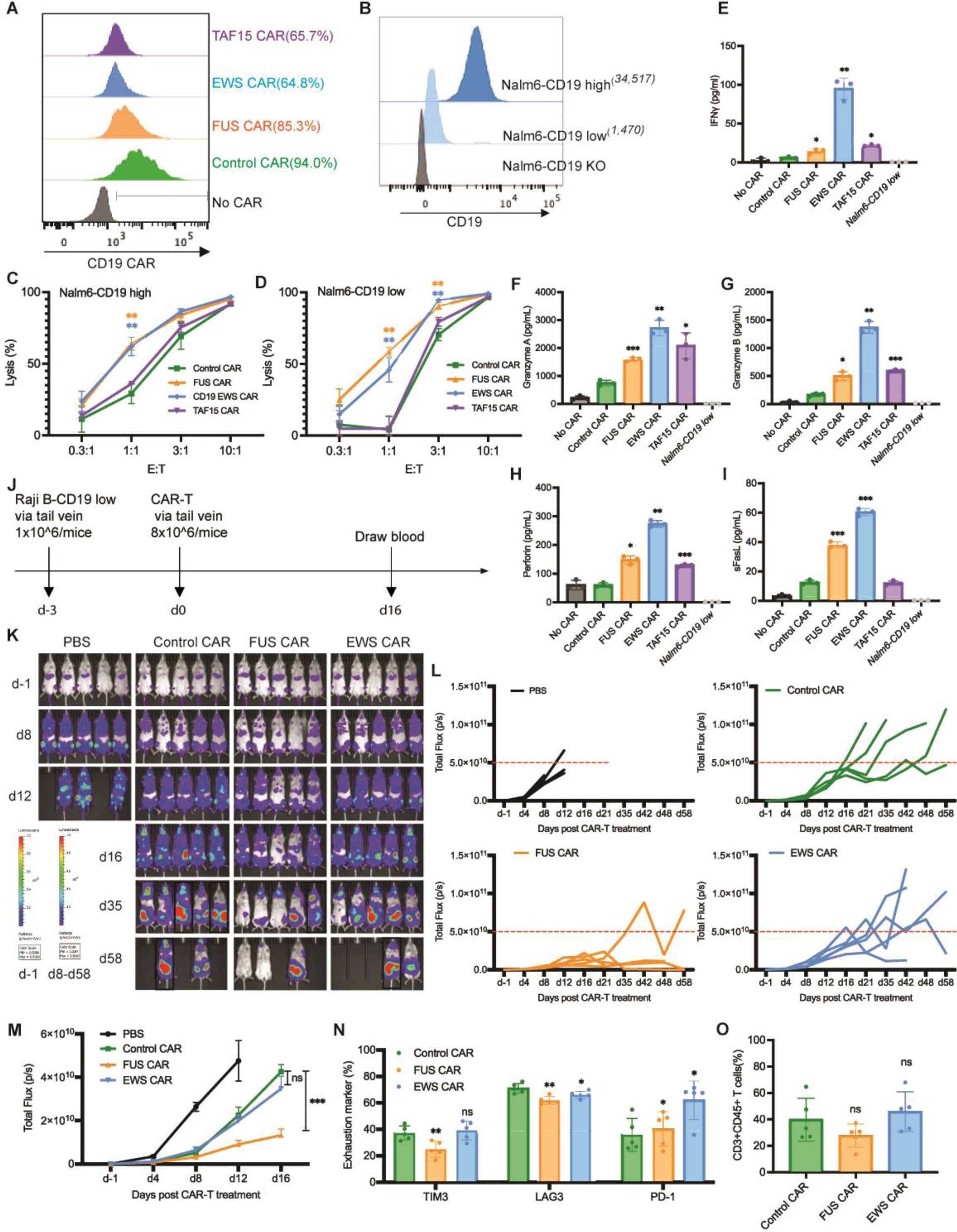
IDR enhanced the cytotoxicity of CD19 CAR-T in vitro and in vivo. A) Expression of the control or IDR CAR targeting CD19 (scFv FMC63) in human primary T cells. CAR expression was measured by flow cytometry using an anti-FMC63 antibody. GFP tag was not included in the IDR CARs because the expression of GFP-tagged IDR CAR decreased over time, but the GFP-tagged control CAR did not. Therefore, to maintain a similar CAR size and expression between the control and IDR CAR over time, the GFP tag was removed from the IDR CAR but maintained in the control CAR. B) Quantification of the CD19 level in Nalm6 cells expressing high (34,517 molecules per cell) or low (1,470 molecules per cell) level of CD19. The average number of CD19 on each cell was determining by flow cytometry using a BD Quantibrite Kit. C-D) Cytotoxicity of CD19 CAR-T in vitro. The control or IDR CAR-Ts were co-cultured with CD19-high or low Nalm6 cells expressing luciferase (Fluc+) for 1 day with an effector to target (E:T) ratio from 0.3:1 to 10:1. The percentile of lysed Naml6 cells were quantified by the luciferase assay and normalized to the group of Nalm6 alone. N=3 independent experiments. Shown are mean ± std (*\*\* p<0.01*). E-I) Production of cytotoxic factors by CAR-T. The control or IDR CAR-Ts were co-cultured with CD19-low Nalm6 cells for 1 day at an E:T of 3:1. The cytokines released into the culture media were quantified by flow cytometry using a LegendPlex assay kit. N=3 independent experiments. Shown are mean ± std (*\* p<0.05, ** p<0.01, *** p<0.001*). J) Examination of the anti-tumor effect of CD19 IDR CAR-Ts in a mouse xenograft model. 1×10^6^ FLuc+ CD19 low Raji B cells were engrafted into the immune-deficient NSG mice via tail vein. Three days later, 8×10^6^ CAR-T cells or PBS were injected into the mice via tail vein. Cancer progression was monitored by bioluminescent imaging. Blood was drawn at Day 16 to characterize T cell phenotypes. K) Tumor progression revealed by bioluminescent imaging. Note that the bioluminescence in d-1 is much smaller than those in other days. It adopted a different heatmap scale in order to display a good dynamic range. Mice in black rectangle displayed a total flux exceeding the limit of end timepoint (1×10^11^ p/s). L-M) Quantification of tumor progression in vivo. Both the individual and averaged traces were shown. N=5 mice. Shown are mean ± std (n.s. p>=0.05, *\*\*\* p<0.001*). N) Expression of T cell exhaustion markers during cancer progression. Blood was collected at day 16 post CAR-T infusion, stained with antibodies recognizing the exhaustion marker TIM3, LAG3, and PD1, and analyzed by flow cytometry. N=5 mice. Shown are mean ± std (n.s. p>=0.05, *\* p<0.05, ** p<0.01*). O) Quantification of T cells number in the blood. Blood was collected at day 16 post CAR-T infusion, stained with antibodies recognizing human CD3 and human CD45, and analyzed by flow cytometry. N=5 mice. Shown are mean ± std (n.s. p>=0.05).

To assess the tumor killing effect of IDR CAR-Ts toward CD19-low cancer cells in vivo, CD19-low Raji B cells expressing a luciferase reporter were injected into the immune-deficient NSG mice intravenously. Three days later, the control, FUS, or EWS CAR-T cells were infused through the tail vein. The cancer progression was monitored by bioluminescence imaging (Fig. 2J). We found that FUS CAR-T inhibited cancer proliferation better than the control CAR-T (Fig. 2K-2M). The expression of T cell exhaustion marker TIM3 and LAG3 were lower in FUS CAR-T (Fig. 2N and S2E). No significant difference was detected in the T cell number (Fig. 2O and S2F) or memory phenotype (Fig. S2G - S2H). Together, these data suggested that FUS CAR-T exhibited an enhanced anti-cancer activity both in vitro and in vivo.

### IDR enhanced cytotoxicity of HER2 CAR-T

To determine if the effect of IDR in promoting cytotoxicity applied to CARs beyond CD19, we constructed IDR CARs targeting HER2 (Fig. 3A), an antigen commonly overexpressed in multiple solid tumors including breast, ovarian, lung and colorectal cancers. Human primary T cells were infected with lentivirus encoding the control, FUS, TAF15, or EWS CAR (Fig. 3B). Similar to the case of CD19 CAR, IDR did not trigger tonic signaling of HER2 CAR as assessed by CD69 expression and TNFα expression (Fig. S1C and S1D). Next, we selected multiple target cell lines for testing cytotoxicity: the lymphoblast K562 cells ectopically expressing high or low HER2 (Fig. 3C), an ovarian cancer cell line SKOV3 expressing high HER2 and a colon cancer cell line HT29 expressing low HER2 (Fig. 3D). These target cells were co-cultured with the control or IDR CAR-Ts. We found that FUS and TAF15 CAR-Ts displayed a higher cytotoxicity towards all four cell lines tested, as compared to the control CAR-T (Fig. 3E-3H). Consistent with that, FUS and TAF15 CAR-Ts secreted a higher level of cytotoxic factors including IFNꝨ, Perforin, and FasL, than the control CAR-T (Fig. 3I – 3M and S3A). Together, these data suggested that FUS and TAF15 enhanced the cytotoxicity of HER2-high and low CAR-Ts in vitro.

**Figure 3.**
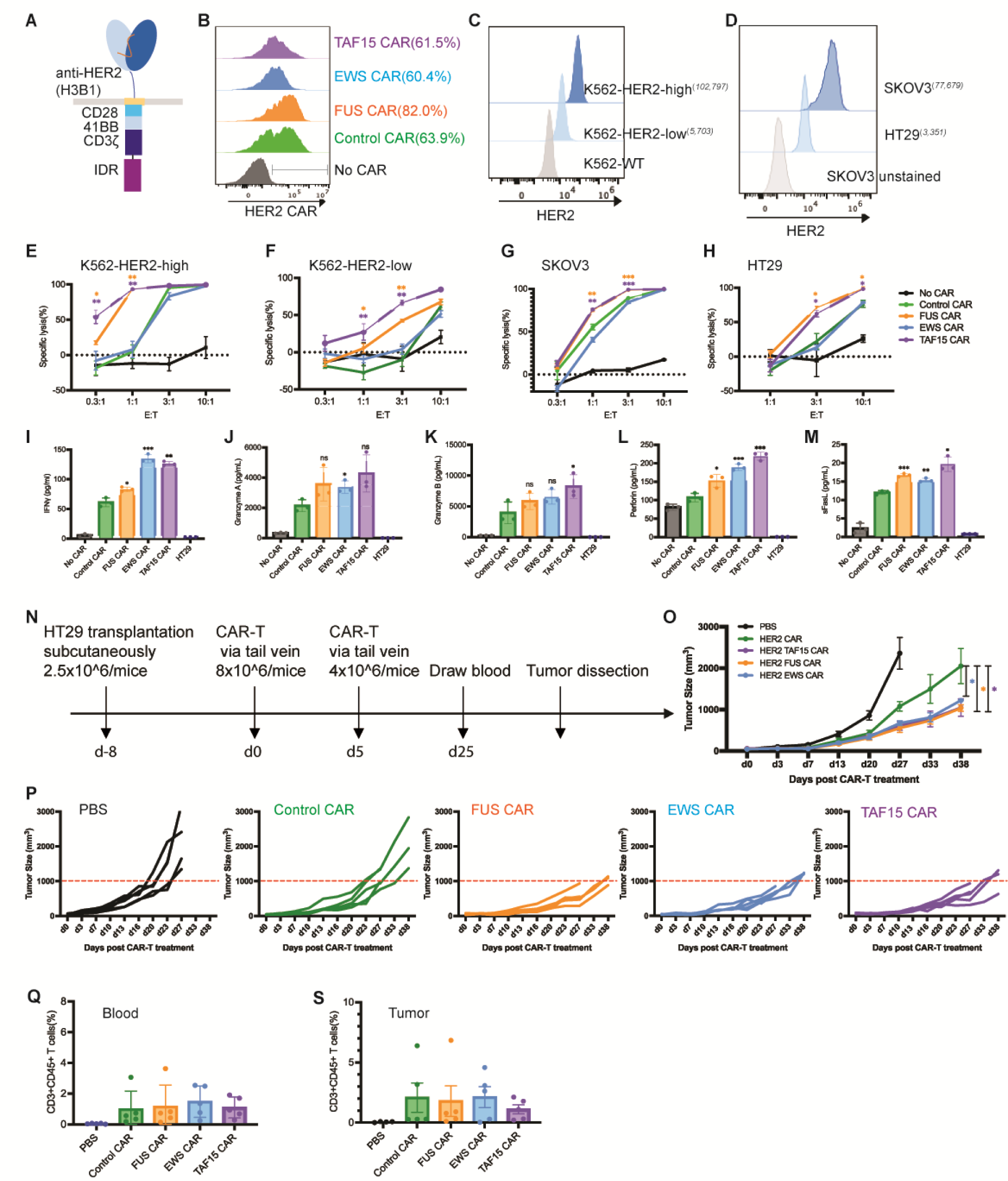
IDR enhanced the cytotoxicity of HER2 CAR-T in vitro and in vivo. A) Schematics of the HER2 CAR used in this study. It contained an scFv targeting HER2 (H3B1), a CD28 hinge and transmembrane domain, an intracellular signaling domain composed of CD28, 41BB and CD3ζ, and an intrinsically disordered region (IDR). The control CAR contained a GFP tag at the C-terminus so that the size and surface expression level of the control and IDR CARs can be maintained in a similar level over time. B) Expression of the control or IDR CAR targeting HER2 in human primary T cells. CAR expression was detected by recombinant HER2 proteins and measured by flow cytometry. C-D) Quantification of the HER2 level in K562 cells expressing high (102,797 molecules per cell) or low (5,703 molecules per cell) HER2, and in SKOV3 (77,679 molecules per cell) and HT29 (3,351 molecules per cell) cells. The average number of HER2 on each cell was determining by flow cytometry using a BD Quantibrite Kit. E-H) Cytotoxicity of HER2 CAR-T in vitro. The control or IDR CAR-Ts were co-cultured with SKOV3, HT29, HER2-high or low K562 cells expressing luciferase (Fluc+) for 1 day with an effector to target (E:T) ratio from 0.3:1 to 10:1. The percentile of lysed SKOV3, HT29, or K562 cells were quantified by the luciferase assay and normalized to the group of SKOV3, HT29, or K562 alone. N=3 independent experiments. Shown are mean ± std (*\* p<0.05, ** p<0.01, *** p<0.001*). I-M) Production of cytotoxic factors by CAR-T. The control or IDR CAR-Ts were co-cultured with HT29 cells for 1 day at an E:T of 3:1. The cytokines released into the culture media were quantified by flow cytometry using a LegendPlex assay kit. N=3 independent experiments. Shown are mean ± std (n.s. p>=0.05, *\* p<0.05, ** p<0.01, *** p<0.001*). N) Examination of the anti-tumor effect of HER2 IDR CAR-Ts in a mouse xenograft model. 2.5×10^6^ HT29 cells were engrafted subcutaneously into the immune-deficient NSG mice. Eight days later, 8×10^6^ CAR-T cells were injected into the mice via tail vein, followed by a second injection of 4×10^6^ CAR-T cells five days after. Cancer progression was monitored by measuring the tumor size using an electronic digital caliper. O-P) Quantification of cancer progression in vivo. Both the averaged and individual traces were shown. N=5 mice. Shown are mean ± std (*\* p<0.05*). Q) Quantification of T cell number in blood. Blood was drawn at Day 25, stained with antibodies recognizing human CD3 and CD45. The percentile of human T cells was measured by flow cytometry. N=5 mice. Shown are mean ± std. R) Quantification of tumor-infiltrating T cells. At the end timepoint (the tumor size over 2000 mm^3^), tumors were dissected, digested, stained with antibodies recognizing human CD3 and CD45. The percentile of human T cells was measured by flow cytometry. N=5 mice. Shown are mean ± std.

To determine the anti-tumor effect of IDR CAR-Ts toward HER2-low cells in vivo, HT-29 cells were injected into the immune-deficient NSG mice subcutaneously. Eight days later, the control or IDR CAR-T cells were infused through the tail vein. The tumor progression was monitored by an electronic caliper over 7 weeks (Fig. 3N). We found that FUS and TAF15 CAR-T inhibited tumor growth better than the control CAR-T (Fig. 3O – 3P), which is consistent with the in vitro killing results. Interestingly, although EWS CAR-T did not show a higher cytotoxicity in vitro, it did display a higher anti-tumor effect than the control CAR-T. This could be potentially explained by the higher cytotoxic factors, including IFNꝨ, that were secreted by EWS CAR-T (Fig. 3I). The numbers of circulating and tumor-infiltrating T cells were comparable between the control and IDR CAR-Ts (Fig.3Q-3S and S3B), suggesting that the enhanced killing was not related to an increased cell number but to an increased cytotoxicity (Fig. 3H). Together, these data suggested that IDR CAR-Ts exhibited an enhanced anti-cancer efficacy towards HER-low cells in vivo.

### IDR enhanced cytotoxicity of a low-signaling CD22 CAR-T

In addition to the anti-CD19 and anti-HER2 CAR which target membrane-proximal epitopes, we also tested how IDRs affect the cytotoxicity of an anti-CD22 CAR (RFB4), which showed very low signaling efficiency due to the membrane-distal epitope position on CD22 (James et al., 2008; Xiao et al., 2022b). Using a similar design strategy to the CD19 CAR, we constructed the IDR CARs targeting CD22 by fusing the IDR on the C-terminus (Fig. 4A). Human primary T cells were infected with lentivirus encoding the control, FUS, EWS, or TAF15 CAR (Fig. 4B). The wild-type Raji B or Nalm6 cells which express a medium level of CD22 (Fig. 4C) were co-cultured with CAR-Ts. We found that FUS and EWS CAR-T displayed a higher cytotoxicity as compared to the control CAR-T when co-cultured with either Raji B or Nalm6 cells (Fig. 4D-4E). We also measured the cytotoxic factors release and found that FUS and EWS CAR-Ts released a slightly higher Granzyme A and B, and FasL than the control CAR-T, though it was not statistically significant (Fig. 4F-4J and S4A). Together, these data showed that FUS and EWS enhanced the cytotoxicity of a low-signaling CD22 CAR-T.

**Figure 4.**
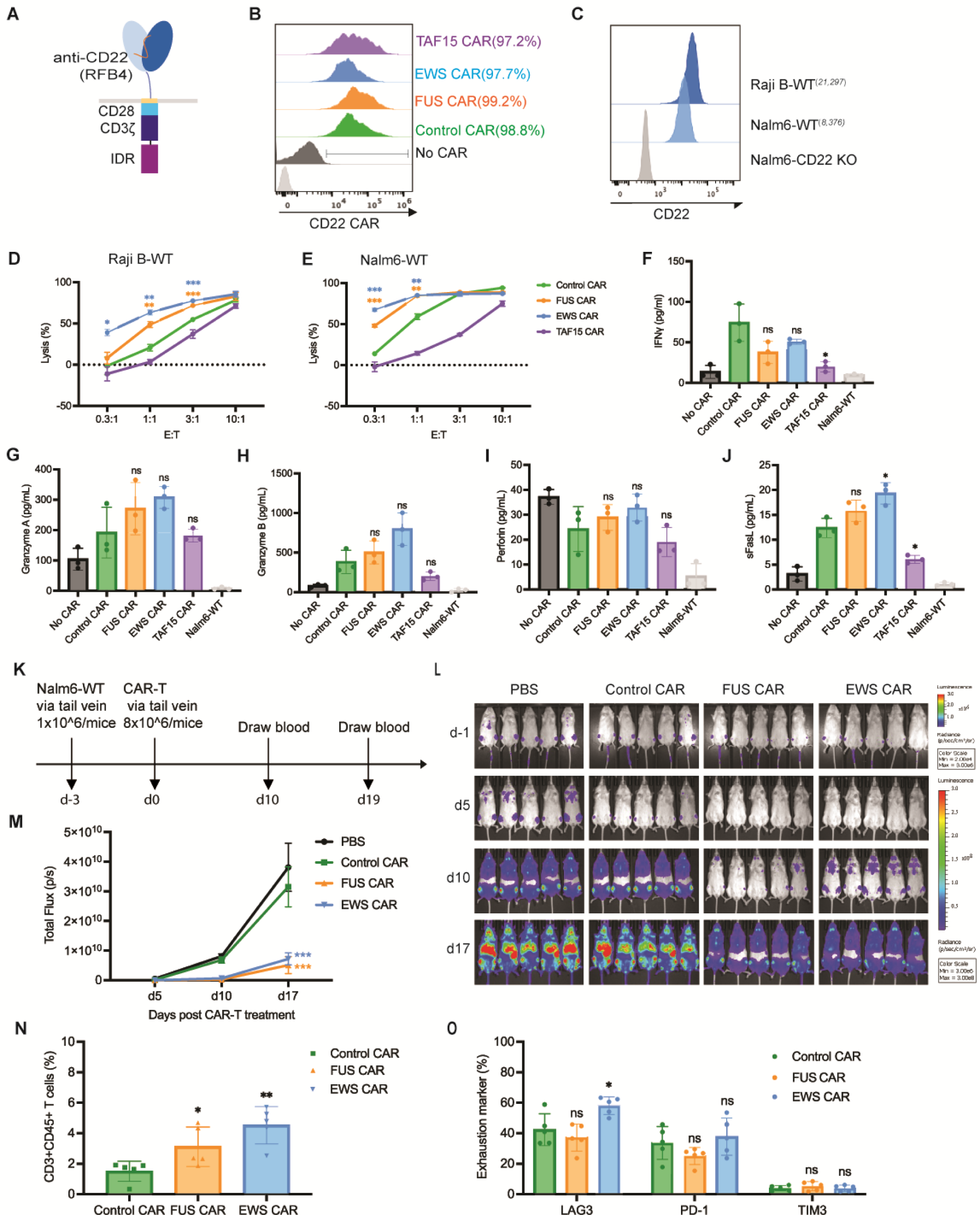
IDR enhanced the cytotoxicity of CD22 CAR-T in vitro and in vivo. A) Schematics of the CD22 CAR used in this study. It contained an scFv targeting CD22 (RFB4, low signaling efficiency), a CD28 hinge and transmembrane domain, an intracellular signaling domain composed of CD28 and CD3ζ, and an intrinsically disordered region (IDR). The control CAR contained a GFP tag at the C-terminus so that the size and surface expression level of the control and IDR CARs were similar. B) Expression of the control or IDR CAR targeting CD22 in human primary T cells. CAR expression was detected by recombinant CD22 proteins and measured by flow cytometry. C) Quantification of the CD22 level in the wild-type Raji B (21,297 molecules per cell) and Nalm6 (8,376 molecules per cell). The average number of CD22 on each cell was determining by flow cytometry using a BD Quantibrite Kit. D-E) Cytotoxicity of CD22 CAR-T in vitro. The control or IDR CAR-Ts were co-cultured with Raji B or Nalm6 cells expressing luciferase (Fluc+) for 3 or 1 day, respectively, with an effector to target (E:T) ratio from 0.3:1 to 10:1. The percentile of lysed Naml6 cells were quantified by the luciferase assay and normalized to the group of Raji B or Nalm6 alone. N=3 independent experiments. Shown are mean ± std (*\* p<0.05, ** p<0.01, *** p<0.001*). F-J) Production of cytotoxic factors by CAR-T. The control or IDR CAR-Ts were co-cultured with the wild-type Nalm6 cells for 1 day at an E:T of 1:1. The cytokines released into the culture media were quantified by flow cytometry using a LegendPlex assay kit. N=3 independent experiments. Shown are mean ± std (n.s. p>=0.05, *\* p<0.05*). K) Examination of the anti-tumor effect of CD22 IDR CAR-Ts in a mouse xenograft model. 1×10^6^ FLuc+ Nalm6-WT cells were engrafted into the immune-deficient NSG mice via tail vein. Three days later, 8×10^6^ CAR-T cells or PBS were injected into the mice via tail vein. Cancer progression was monitored by bioluminescent imaging. Blood was drawn at Day 10 and 19 to characterize T cell phenotypes. L) Tumor progression revealed by bioluminescent imaging. Note that the bioluminescence in d-1 is much smaller than those in other days. It adopted a different heatmap scale in order to display a good dynamic range. M) Quantification of tumor progression in vivo. N=5 mice. Shown are mean ± std (*\*\*\* p<0.001*). N) Quantification of T cells in the blood. Blood was collected at day 10 post CAR-T infusion, stained with antibodies recognizing human CD3 and human CD45, and analyzed by flow cytometry. N=5 mice. Shown are mean ± std (*\* p<0.05, ** p<0.01*). O) Expression of T cell exhaustion markers during cancer progression. Blood was collected at day 19 post CAR-T infusion, stained with antibodies recognizing the exhaustion marker TIM3, LAG3, and PD1, and analyzed by flow cytometry. N=5 mice. Shown are mean ± std (n.s. p>=0.05, *\* p<0.05*).

To determine if IDRs improve the anti-tumor efficacy of the above low-signaling CD22 CAR in vivo, the wild-type Nalm6 cells were injected into the NSG mice intravenously. Three days later, the control, FUS or EWS CAR-T cells were infused through the tail vein. The cancer progression was monitored by bioluminescence imaging (Fig. 4K). Similar to the in vitro killing result, FUS and EWS CAR-T inhibited the tumor growth better than the control CAR-T (Fig. 4L – 4M). The circulating T cell number was higher in the FUS and EWS group than the control group (Fig. 4N and S4B). The expression of exhaustion markers (Fig. 4O and S4C) or memory phenotypes (Fig. S4D and S4E) was not significantly different between the control and IDR CAR-Ts. Together, these data suggested that IDRs from FUS and EWS promote the anti-tumor effect of a low-signaling CD22 CAR.

### IDR enhanced CAR-T activation by promoting CD3ζ and LAT phosphorylation

To determine the mechanism by which IDRs promote CAR-T activation, we monitored membrane-proximal signaling in CAR-Ts upon engagement with CD19. Firstly, we determined the cell-cell conjugation efficiency between the CAR-T and cancer cells as a reflection of binding strength between CAR and antigen. We found that IDRs did not affect the cell-cell conjugation (Fig. 5A), suggesting that IDRs promote CAR-T activation post CAR-antigen interactions. Following a CAR-antigen engagement, the CD3ζ domain on CAR is phosphorylated, which leads a cascade of biochemical reactions occurring at the T cell synapse and intracellular space. We co-cultured CAR-Ts with CD19-low Raji B cells and monitored the antigen-dependent signaling kinetics by flow cytometry. We found that the FUS, EWS, and TAF15 IDR enhanced the phosphorylation of CD3ζ, LAT, ERK, which lined up a pathway that mediates IDR CAR-T activation (Fig. 5B-5D and S5A). Because CD3ζ and LAT are two transmembrane proteins enriched in the CAR-T synapse, we examined their phosphorylation level in the synapse by confocal microscopy. Consistent with the flow cytometry measurement, all three IDRs enhanced the phosphorylation of CD3ζ and LAT in the synapse (Fig. 5E-5H). Together, these data suggested that IDRs promote CAR-T activation against low-antigen cancer cells by promoting membrane-proximal signaling pathways.

**Figure 5.**
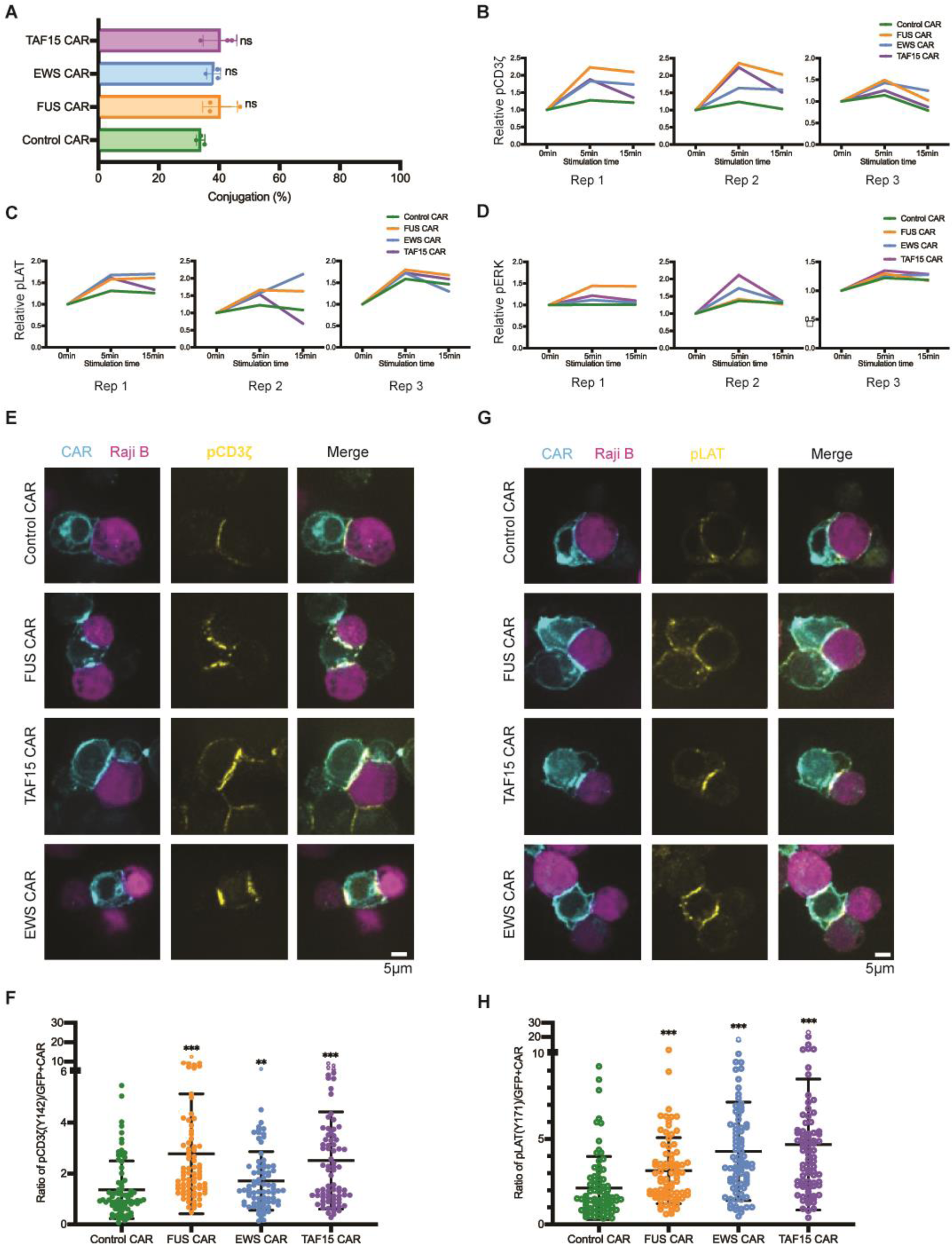
IDR promoted membrane-proximal signaling in CAR-T. A) Cell-cell conjugation between CAR-T and Nalm6 cells. CD19 CAR-Ts were co-cultured with CD19-low RajiB cells for 30 min at 37°C with an E:T ratio of 1:1. Cell conjugation was examined by confocal microscopy. N=3 independent experiments. Shown are mean ± std (n.s. p>=0.05). B-D) Phosphorylation kinetics of CD3ζ (pY142), LAT (pY171), and ERK (pT202/Y204). CAR-Ts were mixed with CD19-low Raji B cells at an E:T=1:1 at 37°C. Cells were fixed at 0, 5 or 15 min, stained and examined with flow cytometry. Displayed are traces for three independent experiments. Relative pCD3z, pLAT, and pERK was calculated by dividing the geometric mean of fluorescence at indicated time point to that at 0 min. E) Phosphorylation of CD3ζ at the CAR-T synapse. CD19 CAR-Ts were co-cultured with CD19-low Raji B cells for 5 min at 37°C with an E:T = 1:1. The cell mixture was fixed and stained with an antibody recognizing CD3ζ (pY142), and examined by confocal microscopy. F) Quantification of pCD3ζ at the CAR-T synapse. Normalized pCD3ζ was calculated by dividing the fluorescence of pCD3ζ to the fluorescence of CAR in the synapse. N=77-107 cells. Shown are mean ± std (*\*\* p<0.01, *** p<0.001*). G) Phosphorylation of LAT at the CAR-T synapse. CD19 CAR-Ts were co-cultured with CD19-low Raji B cells for 5 min at 37°C with an E:T = 1:1. The cell mixture was fixed and stained with an antibody recognizing LAT (pY171), and examined by confocal microscopy. H) Quantification of pLAT at the CAR-T synapse. Normalized pLAT was calculated by dividing the fluorescence of pLAT to the fluorescence of CAR in the synapse. N= 79-105 cells. Shown are mean ± std (*\*\*\* p<0.001*).

### Oligomerization of CAR by coiled-coil domains caused reduced surface localization

In addition to IDR-mediated protein condensation, the coiled-coil domain is a commonly used tool to induce oligomerization of protein of interest. To test if coiled-coil domains could promote CAR-T activation, we fused coiled-coil (cc) domains that mediate dimerization, tetramerization, or hexamerization (Deng et al., 2006; Spencer and Hochbaum, 2017) to the C-terminus of a CD19 CAR (Fig. 6A) and introduced these cc-CARs into human primary T cells. Interestingly, whereas the cc-dimer maintained a similar cell surface expression level as compared to the control CAR, cc-tetramer and hexamer CAR showed a dramatic reduction in the cell surface localization (Fig. 6B). Consequently, the cell-cell conjugation percentage between CAR-T and Nalm6 cells was significantly reduced in the cc-tetramer and hexamer as compared to the control CAR (Fig. 6C). In consistency, CAR-T activation, as evaluated by CD69 expression (Fig. 6D and S6A) and IFNꝨ secretion (Fig. 6E), was significantly reduced in cc-tetramer or hexamer. The reduction in cell-cell conjugation and CAR-T activation were recapitulated when using Raji B as a target cell (Fig. S6B to S6D). Similar to CAR, the oligomerization-induced receptor internalization was frequently observed in other transmembrane receptors including EGFR and FGFR (Hofman et al., 2010; Pozniak et al., 2020). The fact that IDRs do not affect cell surface expression of CAR suggested that IDRs present a unique advantage to promote CAR clustering without causing enhanced receptor internalization.

**Figure 6.**
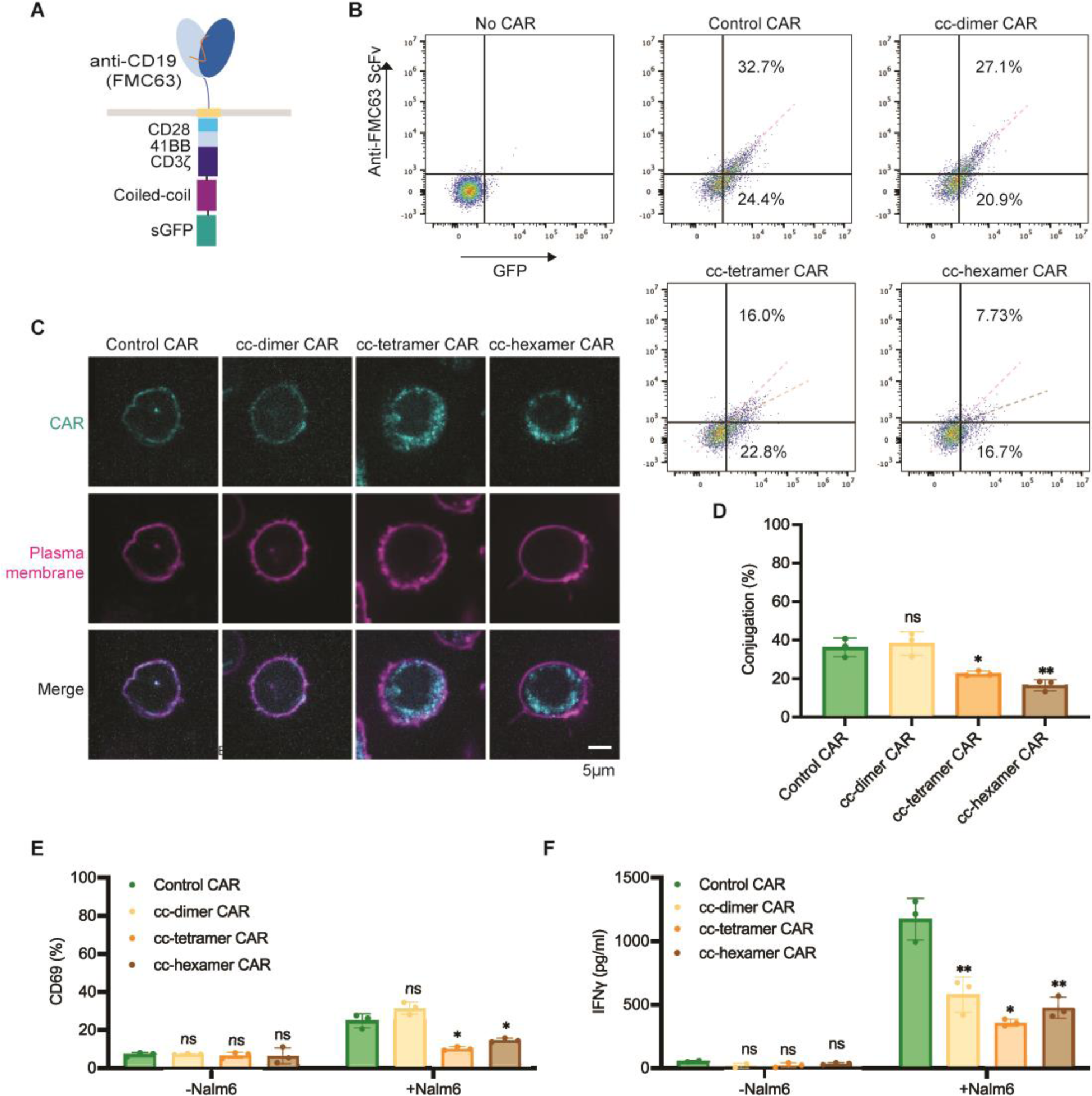
Oligomerization by coiled-coil domain reduced CAR surface localization and CAR-T activation. A) Schematics of the coiled-coil (cc) CD19 CAR. It contained an scFv targeting CD19 (FMC63), a CD8 hinge, a LAT transmembrane domain, an intracellular signaling domain composed of CD28, 41BB, and CD3ζ, and a coiled-coil domain. B) Expression of coiled-coil CARs. The localization of CAR on the T cell surface was detected via flow cytometry using an anti-FMC63 antibody. A GFP tag was fused on the C-terminus of each CAR to monitor the total expression of CAR. C) Confocal microscopy revealed the cellular localization of coiled-coil CAR. The plasma membrane was labeled by a CellMask Deep Red dye. D) Cell-cell conjugation between CAR-T and Nalm6 cells. CAR-Ts were mixed with Naml6-H8 clone (expressing 137,795 CD19 molecules per cell) for 30 min at 37°C with an E:T =1:1. Cell conjugation was monitored by confocal microscopy. N=3 independent experiments. Shown are mean ± std (n.s. p>=0.05, *\* p<0.05, **p<0.01*). E-F) Activation of coiled-coil CAR-Ts assessed by CD69 expression and IFNꝨ release. CAR-Ts were co-cultured with Nalm6-H8 for 1 day at an E:T = 1:1. CD69 expression was revealed by flow cytometry. IFNꝨ release was quantified by ELISA. N=3 independent experiments. Shown are mean ± std (n.s. p>=0.05, *\* p<0.05, **p<0.01*).

## Discussion

Biomolecular condensation has been demonstrated to regulate diverse cellular signaling processes (Case et al., 2019; Li et al., 2012; Li et al., 2022). Previous work showed that condensation of TCR signaling molecules promotes membrane-proximal signaling including LAT phosphorylation, RAS activation, actin polymerization, and ERK activation (Huang et al., 2019; Su et al., 2016; Zeng et al., 2021). Our work identified IDR-induced condensation as a new strategy to improve the function of CAR-T. We demonstrated the proof-of-concept application of implementing IDRs to enhance the cytotoxicity of CAR-Ts without inducing tonic signaling. Because IDRs are present in 30% of the human proteins, it is likely that we only revealed a tip of iceberg of the IDRs that can be used for engineering CAR; many more IDRS are expected to be further revealed to modulate the condensation, antigen-binding, and conformation of CARs.

In this work, we demonstrated that several well-characterized IDRs including those from FUS, EWS, and TAF15, promoted the condensation of CARs on the T cell membranes, which leads to an enhanced cytotoxicity towards low antigen-expressing cancer cells. Because the three above IDR-containing proteins are naturally functioning in the nucleus and none of these IDRs have been reported to have a function in T cell signaling or cytotoxicity, our strategy can serve as an orthogonal way to enhance the low signaling efficiency of CAR-T, in combination with other strategies, including selecting specific transmembrane or co-signaling domains (Heitzeneder et al., 2022; Majzner et al., 2020; Priceman et al., 2018), adding new signaling binding motifs (Salter et al., 2021), replacing the intracellular part with that from TCR (HIT) (Mansilla-Soto et al., 2022), dual targeting by CAR together with chimeric costimulatory receptors (Katsarou et al., 2021), or implementing other signaling pathways (Wilkens et al., 2022), to achieve synergistic effects.

It is noted that not all the IDRs tested in our studies promoted CAR condensation. We showed that FUS, EWS, or TAF15 promoted CAR condensation, but not NUP98, TDP43 or SynIDR. This could be attributed to the fact that the ability of these IDRs to promote condensation was initially characterized in the 3D cytosolic environment, which is different from the 2D plasma membrane environment where CAR is localized. The local molecular crowdedness or the negative charge in the inner leaflet of plasma membrane could influence the ability of individual IDRs to promote CAR condensation.

Our work showed that IDRs promoted LAT phosphorylation in the CAR synapse. LAT is one of the key adaptor proteins in the TCR pathway; it forms signaling condensates, recruits and activates a plethora downstream effector to amply TCR signaling (Bunnell et al., 2002; Zhang et al., 1998). Previous studies suggested that LAT is poorly activated (phosphorylated) following CAR-antigen interactions (Dong et al., 2020; Salter et al., 2021), which could explain the low antigen sensitivity of CAR signaling. Here we demonstrated that IDRs can significantly increase the activation of LAT and downstream pathways including SLP76 and ERK, which finally lead to an enhanced cytotoxicity.

Last but not the least, we revealed an interesting difference between the outcomes of different ways to induce CAR clusters. IDR-induced CAR condensation enhanced CAR phosphorylation and CAR-T activation whereas coiled-coil domains reduced cell surface expression of CAR and suppressed CAR-T activation. One of the major differences between IDR and coiled-coil is that IDRs promote self-assembly of CAR through weak interactions and the condensates remain liquid-like or gel-like properties. In contrast, coiled-coil domains induced stable oligomerization which could either affect the trafficking of CAR to the cell surface or trigger instant receptor internalization. Our work suggested that IDRs present a unique advantage in promoting the assembly of CAR into a higher order structure without compromising their cell surface localization.

## Acknowledgement

We thank the R. Majzner Lab for sharing CD19-high and low Nalm6 cell lines, S. Chen Lab for sharing HT29 and Nalm6 cell lines, J. Lu Lab for sharing K562 cell lines, E. Ratner Lab for sharing SKOV3 cell lines, and C. Burd for providing comments on the manuscript. Funding: X.S. was supported by an American Cancer Society Research Scholar Grant, the Charles H. Hood Foundation Child Health Research Awards, the Andrew McDonough B+ Foundation Research Grant, the Gilead Sciences Research Scholars Program in Hematology/Oncology, the Rally Foundation A Collaborative Pediatric Cancer Research Awards Program, the Yale SPORE in skin cancer DRP Award CA121974, the Yale Cancer Center Pilot Award, the Yale DeLuca Pilot Award, the NIGMS MIRA (R35) program GM138299, the Gabrielle’s Angel Foundation Medical Research Award, the Pershing Square Sohn Prize for Young Investigators in Cancer research, and the Human Frontier Science Program Early-Career Research Grant. X.Z. was supported by the Leslie Warner Postdoctoral Fellowship. Longhui Zeng was supported by the CRI-Irvington Postdoctoral Fellowship. Fawzaan Hashmi was supported by the Yale College Dean’s Summer Research Fellowship.

## Author contributions

The project was conceptualized by X.Z. and X.S. X.Z., Q.X., L.Z., and F.H. conducted experiments and performed data analysis. Funding supporting this project was obtained by X.S. X.S. supervised the project. The manuscript was written by X.Z. and X.S.

---

**Figure S1.**
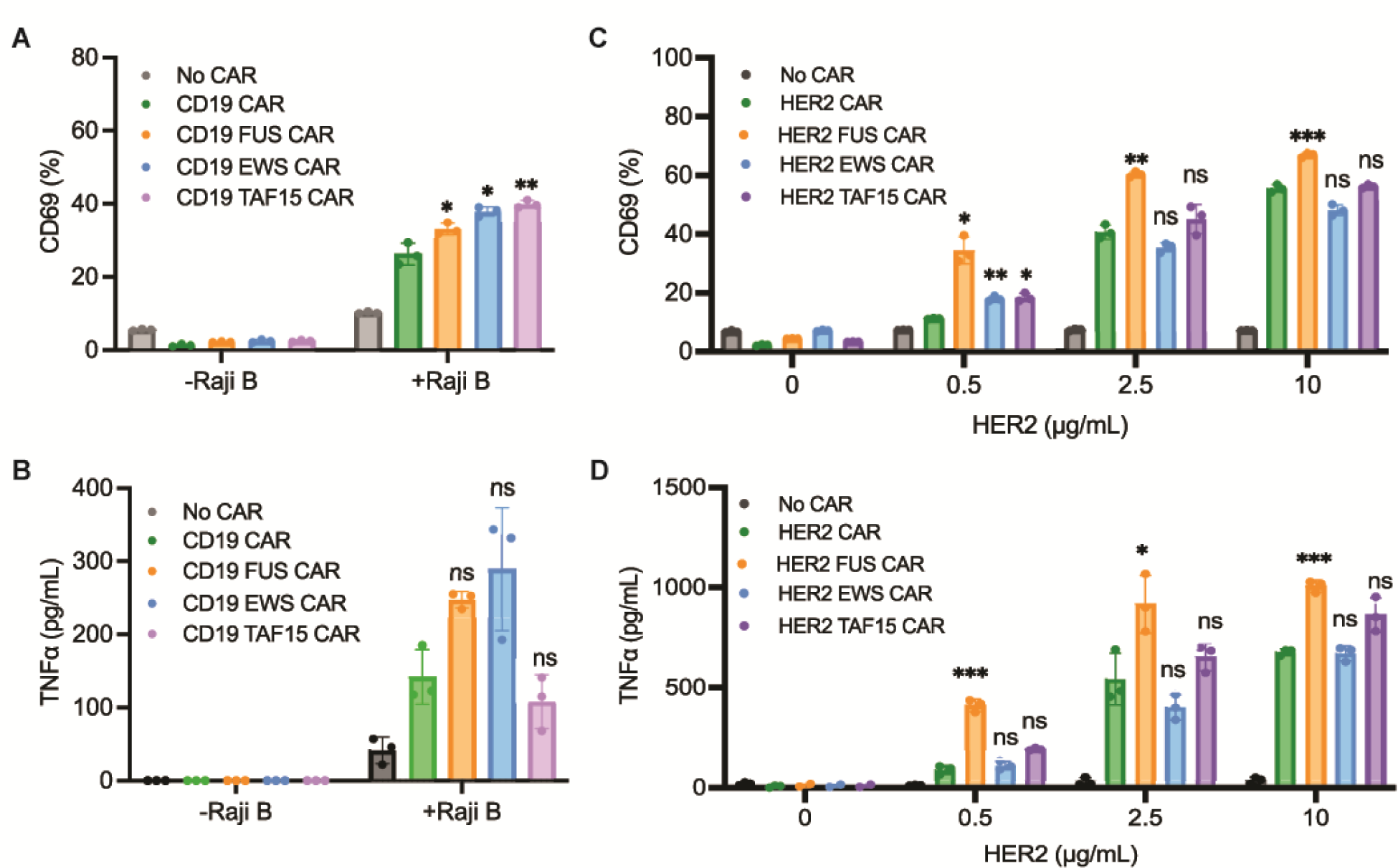
IDR did not trigger tonic signaling. A-B) Activation of CD19 CAR-T in the presence and absence of antigen. CAR-Ts were cultured alone or co-cultured with the wild-type Raji B cells at E:T = 10:1 for 1 day. Expression of CD69 was examined by flow cytometry. TNFα release was measured by ELISA. N=3 independent experiments. Shown are mean ± std (n.s. p>=0.05, *\* p<0.05, **p<0.01*). C-D) Activation of HER2 CAR-T in the presence and absence of antigen. CAR-Ts were stimulated by plate-coated HER2 protein for 1 day. Expression of CD69 was examined by flow cytometry. TNFα release was measured by ELISA. N=3 independent experiments. Shown are mean ± std (n.s. p>=0.05, *\* p<0.05, **p<0.01, ***p<0.001*).

**Figure S2.**
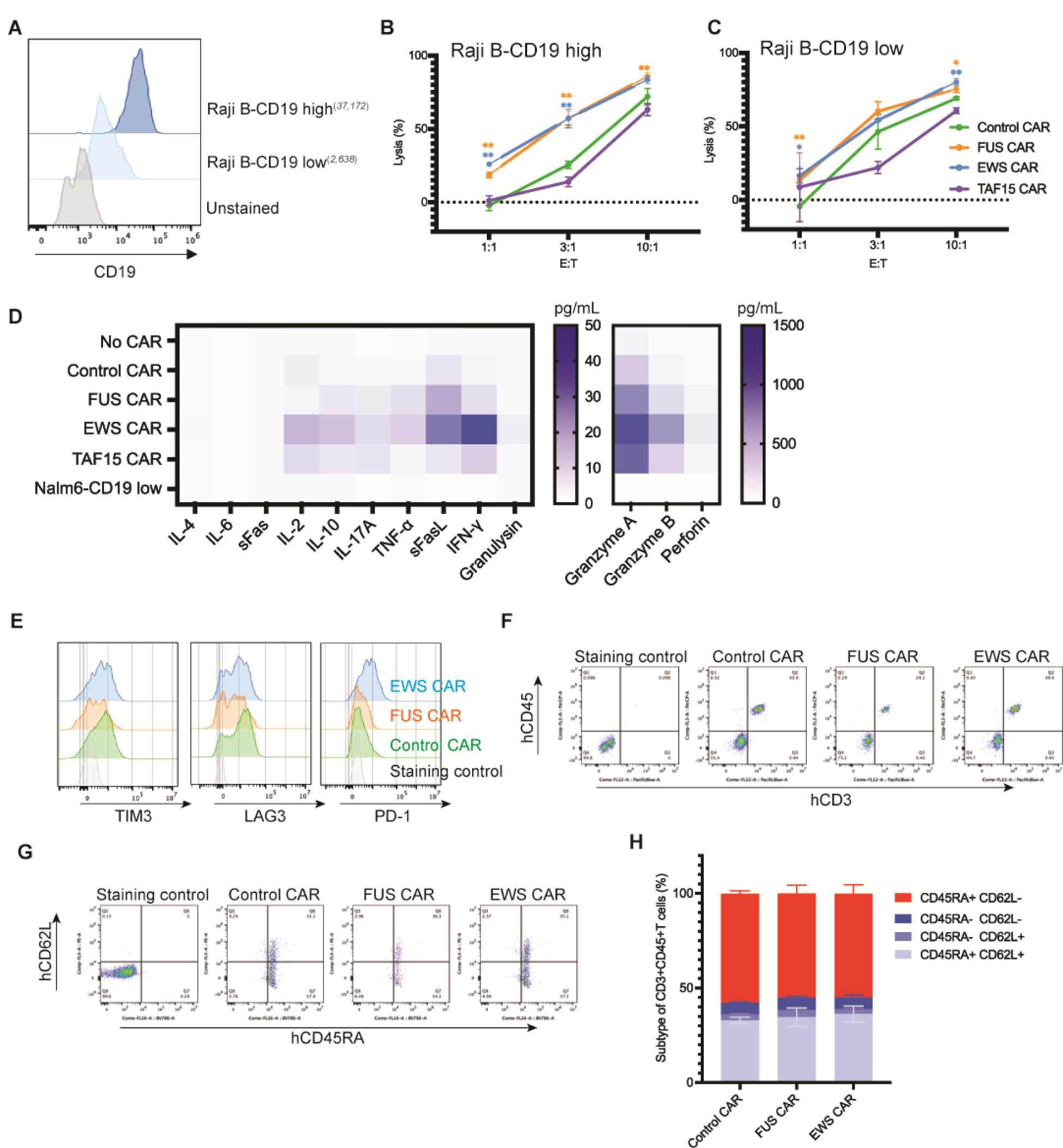
Characterization of IDR CAR-Ts targeting CD19. A) Quantification of the CD19 level in Raji B cells expressing high (37,172 molecules per cell) or low (2,638 molecules per cell) level of CD19. The average number of CD19 on each cell was determining by flow cytometry using a BD Quantibrite Kit. B-C) Cytotoxicity of CD19 CAR-T against Raji B cells. The control or IDR CAR-Ts were co-cultured with CD19-high or low Raji B cells expressing luciferase (Fluc+) for 3 days with an effector to target (E:T) ratio from 1:1 to 10:1. The percentile of lysed Raji B cells were quantified by the luciferase assay and normalized to the group of Raji B alone. N=3 independent experiments. Shown are mean ± std (\**p<0.05, ** p<0.01*). D) Representative heat map illustrating cytotoxic factors released by CD19 CAR-Ts. The control or IDR CAR-Ts were co-cultured with CD19-low Nalm6 cells for 1 day at an E:T of 3:1. The cytokines released into the culture media were quantified by flow cytometry using a LegendPlex assay kit. E) Representative flow cytometry plot showing T cell exhaustion markers. Blood was collected at day 16 post CAR-T infusion, stained with antibodies recognizing the exhaustion marker TIM3, LAG3, and PD1, and analyzed by flow cytometry. F) Representative flow cytometry plot showing T cell population in the blood. Blood was collected at day 16 post CAR-T infusion, stained with antibodies recognizing human CD3 and human CD45. G) Representative flow cytometry plot showing T cell differentiation in vivo. Blood was collected at day 16 post CAR-T infusion, stained with antibodies recognizing CD45RA and CD62L, and analyzed by flow cytometry. H) Quantification of T cell differentiation in vivo. N=5 mice. Shown are mean ± std.

**Figure S3.**
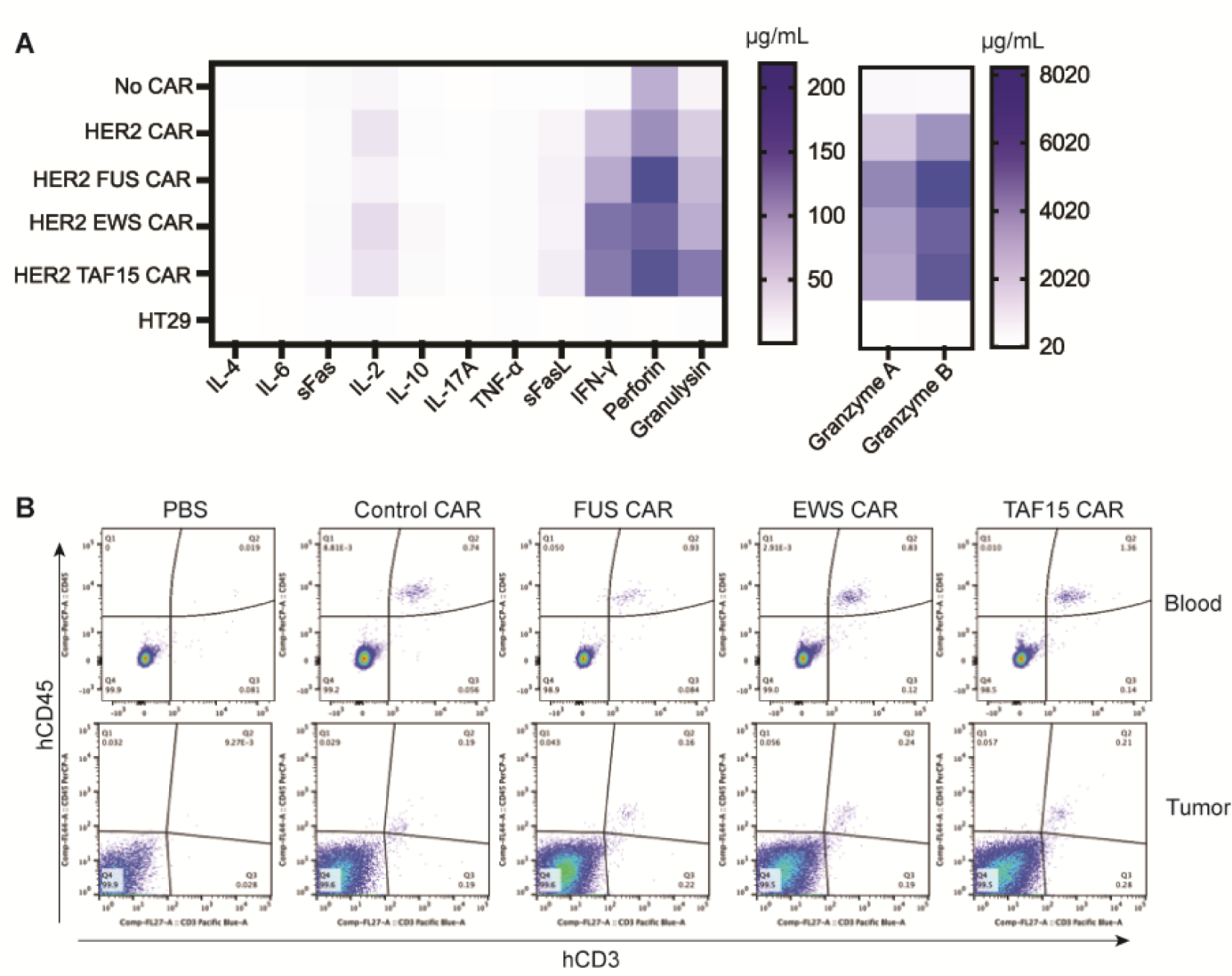
Characterization of IDR CAR-Ts targeting HER2. A) Representative heat map illustrating cytotoxic factors released by HER2 CAR-Ts. The control or IDR CAR-Ts were co-cultured with HT29 cells for 1 day at an E:T of 3:1. The cytokines released into the culture media were quantified by flow cytometry using a LegendPlex assay kit. B) Representative flow cytometry plot showing T cell population in the blood and tumor. Blood was draw at day 25 and tumors were dissected at day 48, digested, stained with antibodies recognizing human CD3 and CD45. The percentile of human T cells was measured by flow cytometry.

**Figure S4.**
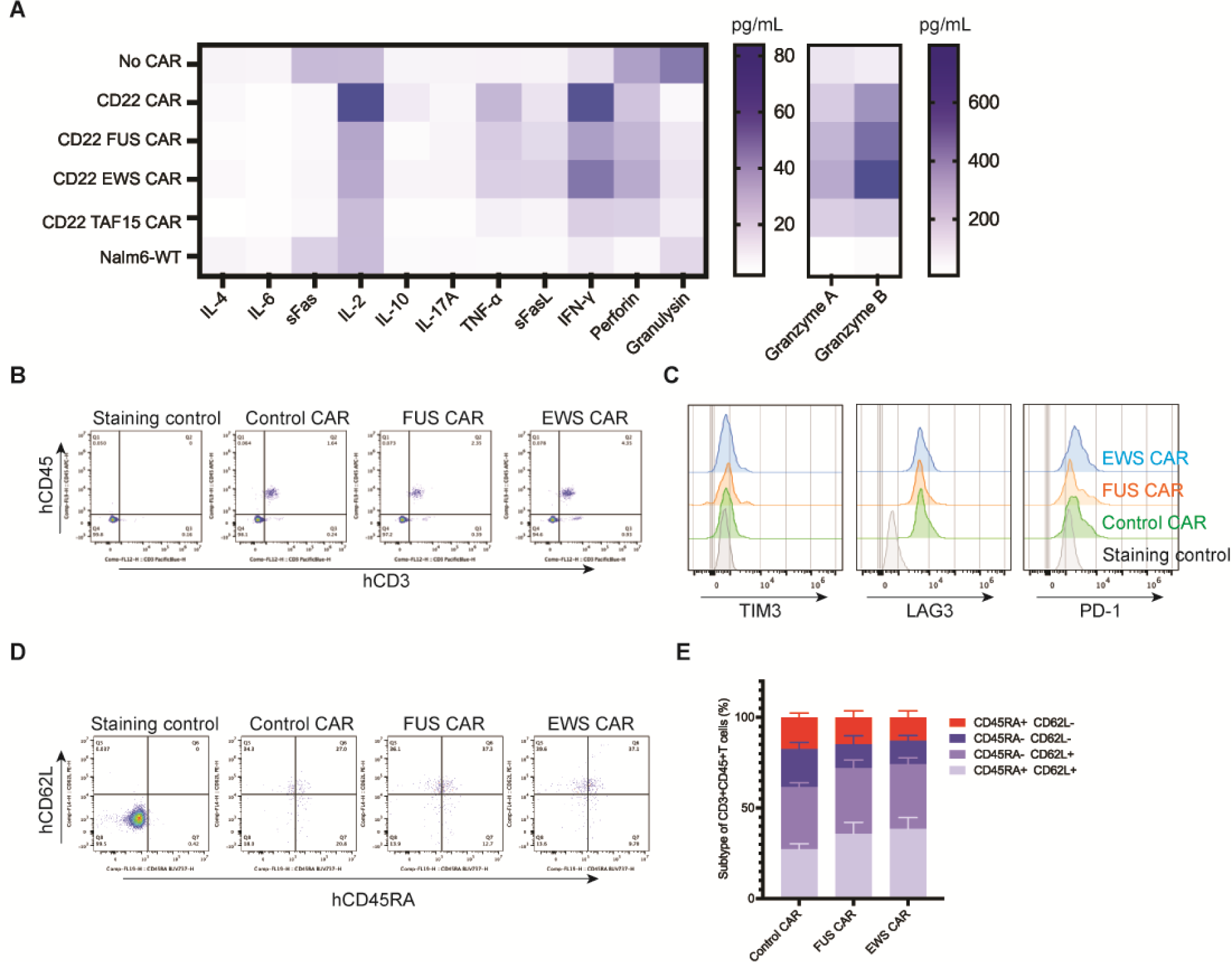
Characterization of IDR CAR-Ts targeting CD22. A) Representative heat map illustrating cytotoxic factors released by CD22 CAR-Ts. The control or IDR CAR-Ts were co-cultured with the wild-type Nalm6 cells for 1 day at an E:T of 1:1. The cytokines released into the culture media were quantified by flow cytometry using a LegendPlex assay kit. B) Representative flow cytometry plot showing T cell population in the blood. Blood was collected at day 10 post CAR-T infusion, stained with antibodies recognizing human CD3 and human CD45. F) Representative flow cytometry plot showing T cell exhaustion markers. Blood was collected at day 19 post CAR-T infusion, stained with antibodies recognizing the exhaustion marker TIM3, LAG3, and PD1, and analyzed by flow cytometry. G) Representative flow cytometry plot showing T cell differentiation in vivo. Blood was collected at day 10 post CAR-T infusion, stained with antibodies recognizing CD45RA and CD62L, and analyzed by flow cytometry. H) Quantification of T cell differentiation in vivo. N=5 mice. Shown are mean ± std.

**Figure S5.**
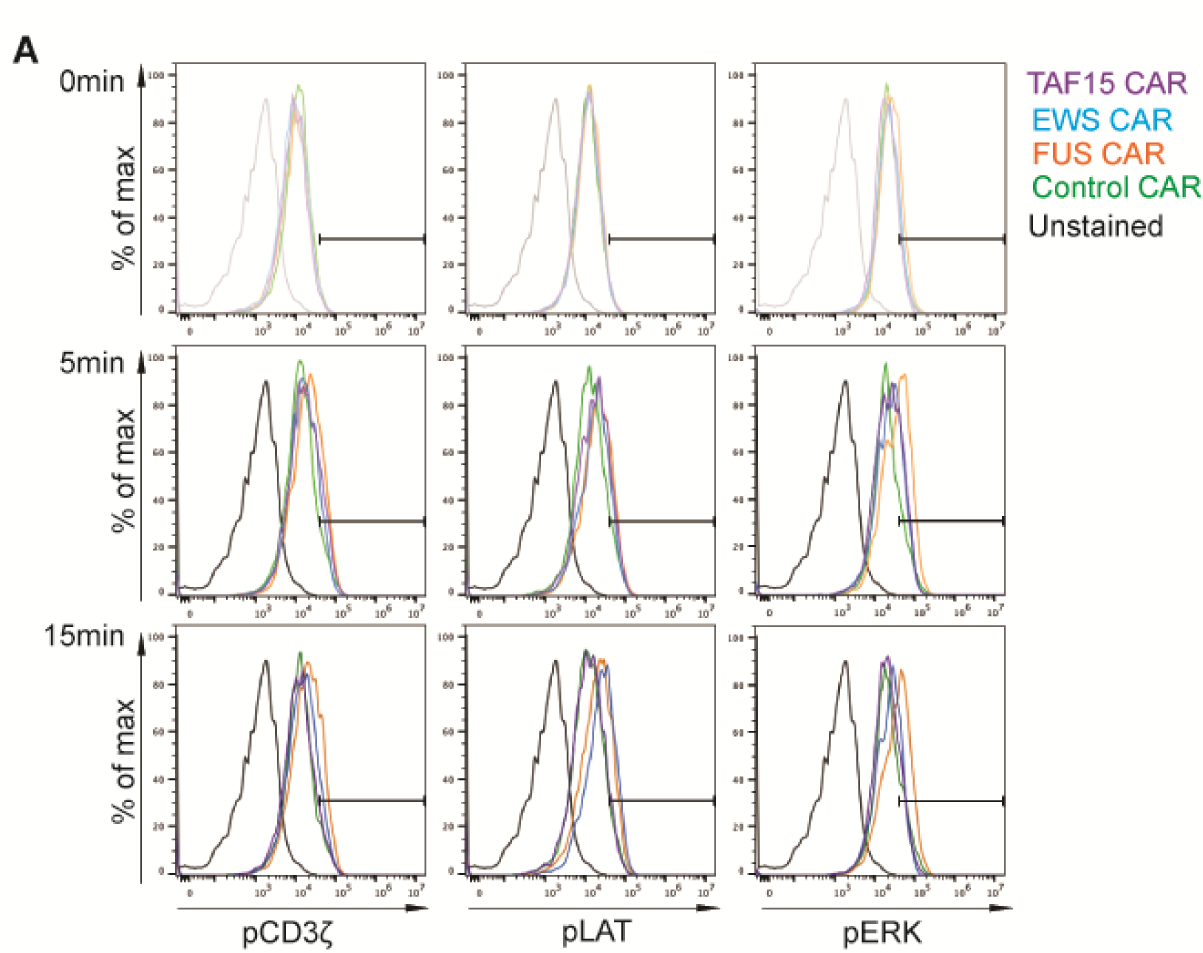
Phosphorylation kinetics of CD3ζ, LAT, and ERK. A) CAR-Ts were mixed with CD19-low Raji B cells at an E:T=1:1 at 37°C. Cells were fixed at 0, 5 or 15 min, stained with phospho-antibodies and examined with flow cytometry. Show are representative flow plots.

**Figure S6.**
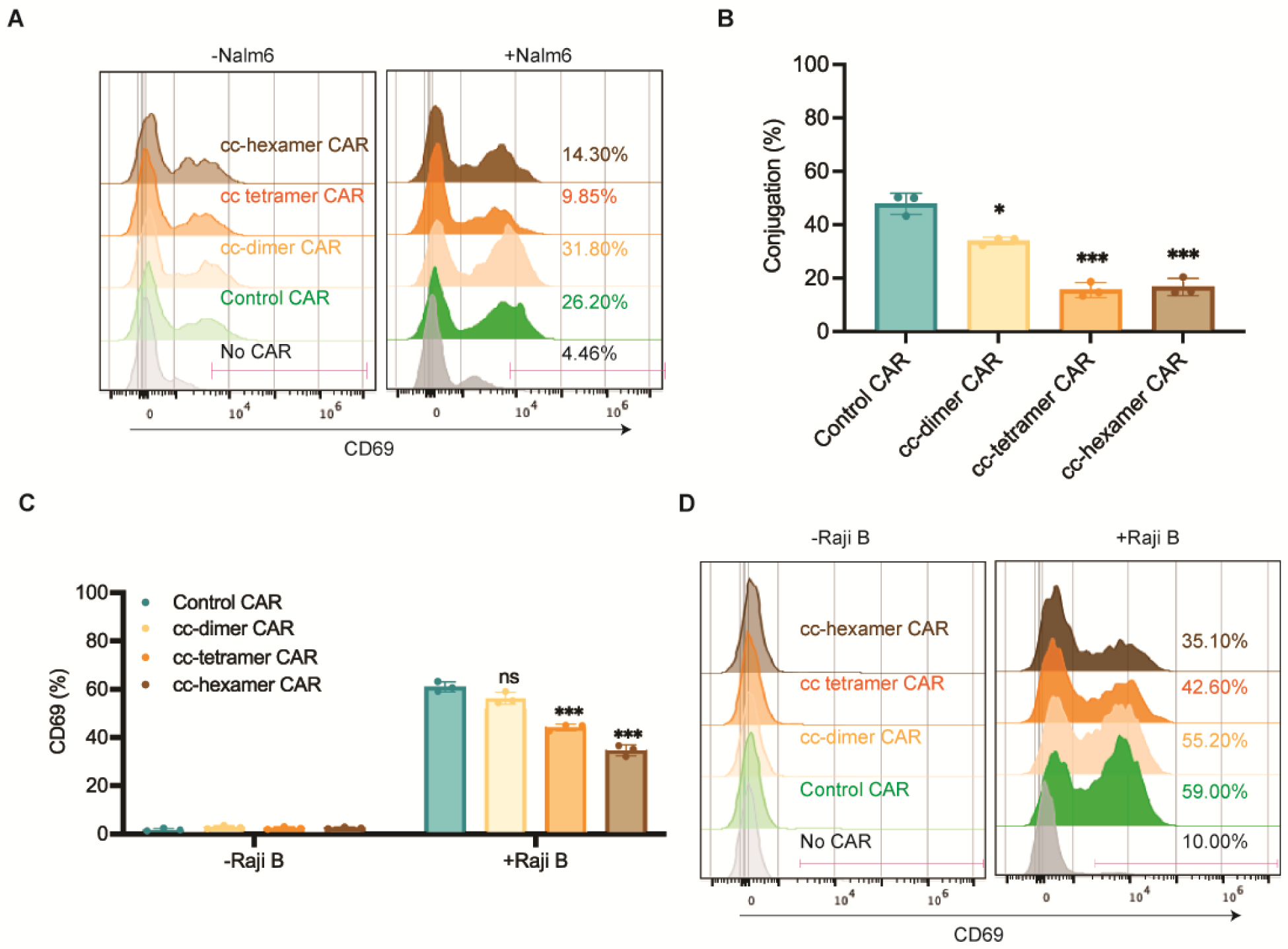
Oligomerization by coiled-coil domain reduced CAR-T activation. A) Representative flow cytometry plots showing CD69 expression in coiled-coil CAR-Ts. CAR-Ts were co-cultured with Nalm6-H8 for 1 day at an E:T = 1:1. B) Cell-cell conjugation between CAR-T and Raji B cells. CAR-Ts were mixed with CD19-high Raji B cells for 30 min at 37°C with an E:T =10:1. Cell conjugation was monitored by confocal microscopy. N=3 independent experiments. Shown are mean ± std (*\* p<0.05, ***p<0.001*). C-D) CD69 expression on coiled-coil CAR-Ts when engaged with Raji B cells. CAR-Ts were co-cultured with CD19-high Raji B cells for 1 day at an E:T = 10:1. CD69 expression was revealed by flow cytometry. N=3 independent experiments. Shown on are mean ± std (n.s. p>=0.05, *\*\*\*p<0.001*). Representative flow cytometry plots were displayed.

## Materials and Methods

### 1. Plasmids and lentivirus

DNA fragments encoding CAR or antigen molecules was inserted into a pHR lentiviral vector with an SFFV promoter and a WPRE terminator. HEK293T cells, maintained in DMEM medium supplemented with 10% FBS and a Glutamine–Penicillin–Streptomycin mix, were co-transfected with the pHR plasmids and second-generation lentiviral packaging plasmids pMD2.G and psPAX2 (Addgene plasmid #12259 and #12260) using Genejuice transfection reagent (EMD Millipore, #70967-3). 48 hrs after transfection, cell culture media containing viral particles were harvested, centrifuged, and filtered through 0.45 μm pore size filters.

### 2. Generation of CAR-T cells

Pan T cells were isolated from PBMCs (Zenbio #SER-PBMC-200) from healthy donors using EasySep™ Human T Cell Isolation Kit (Stem Cell Cat#17951). T cell proliferation was stimulated with Human T-Activator CD3/CD28 Dynabeads (ThermoFisher, #11161D). Cells were cultured in RPMI1640 supplemented with 10% FBS, 50 nM 2-mercaptoethanol, 300 U/mL Interleukin-2 (PEPROTECH, # 200-02). Two days after stimulation, the cells were infected with fresh lentivirus via spinoculation at 800x g for 90 min at 32 °C. Half of the cell culture media were changed with fresh T cell culture medium 24 hours after infection. Five days after infection, Dynabeads were removed, and CAR-T cells were resuspended in a fresh culture medium. The medium was exchanged every two days after.

### 3. Generation of cancer cell lines

The Nalm6, Raji B, and K562 cells were cultured in RPMI1640 supplemented with 10% FBS and a Glutamine–Penicillin–Streptomycin mix. HT29 cells were cultured in DMEM supplemented with 10% FBS and a Glutamine–Penicillin–Streptomycin mix. The Nalm6 CD19-high and -low cell lines expressing GFP and Luciferase were kindly provided by the Majzner Lab at Stanford University. The Raji B CD19-high and -low cells were generated by sorting the wild-type Raji B cells for high and low CD19 expression level. The K562 HER2-high and -low cells were generated by infecting K562 cellls with lentivirus encoding the wild-type HER2, followed by single-cell sorting. The number of antigen molecules per cell was quantified using the BD Quantibrite PE Phycoerythrin Fluorescence Quantitation Kit (BD Bioscience, Cat #340495) by flow cytometry. Luciferase-expressing cells were generated by infecting cancer cells with a lentiviral plasmid expressing Luciferase-GFP or Luciferase-mCherry, followed by FACS to generate stable cell lines.

### 4. *In vitro* cytotoxicity assay

Cytotoxicity was measured by luciferase assay. Cancer cells expressing luciferase were resuspended in RPMI medium supplemented with 10% FBS and mixed with CAR-T cells at an effector to target ratio from 0.3:1 to 10:1. After 24 hours of incubation at 37°C, cells were collected and washed with PBS. Cell pellets were lysed in a lysis reagent (Promega, Cat# 1531). The luminescence of lysates was detected by Luciferase Assay System (Promega, Cat# E1500) and analyzed using a plate spectrophotometer. The spontaneous release control was set up using cancer cells alone. Cell lysis% = [1-(experimental readout-spontaneous readout) /spontaneous readout]x100.

### 5. Cytokine production

CAR-T cells and cancer cells were cocultured at indicated E:T ratios for 24 hours at 37°C. The supernatant was collected for cytokine measurement using ELISA kits (IL2 ELISA kit, BioLegend #431801; IFNγ ELISA kit, BioLegend #430101; TNFα ELISA kit, BioLegend # 430204) or using LEGENDplex™ Human CD8/NK Panel Kit (Cat# 741186), according to the manufacturer’s instruction.

### 6. Flow Cytometry

To determine the cell-surface expression, cells were collected and blocked with an anti-human Fc receptor binding inhibitory antibody in the staining buffer (PBS with 2% FBS and 1 mM EDTA) for 15 minutes at 4°C, which were further incubated with individual antibodies in the staining buffer for 30 minutes on ice. The stained cells were washed twice with the staining buffer before sending for flow cytometry analysis. To determine the intracellular expression of targets of interest, cells were collected and fixed with Fixation/Permeabilization Solution (Cat# 554714) for 15 min on ice, blocked in the staining buffer for 30 min on ice, followed by antibody staining. To characterize T cells in the mice blood, blood was drawn from a tail cut and diluted into PBS supplemented with 3 mM EDTA. Red blood cells were lysed with a red blood cell lysis buffer. The rest of cells were stained as described above and followed by flow cytometry. To characterize tumor-infiltrating T cells, tumors were dissected and digested with RPMI medium containing 0.5 mg/mL Collagenase P and 1 μg/mL DNase per 100 mg of tumor tissues for 30 min at 37°C on a shaker. The digested tumor tissues were further homogenized and passed through a 40 μm strainer, followed by centrifugation to collect cell samples. Cells were further stained and analyzed by flow cytometry as described above.

### 7. Mouse tumor xenograft

Mice were housed in pathogen-free conditions and cared for in accordance with US National Institutes of Health guidelines, and all procedures were approved by the Yale University Animal Care and Use Committee. To generate liquid cancer models, 1 million of Raji B or Naml6 cells in 100 uL of PBS were intravenously injected into the NSG (NOD.Cg-Prkdcscid Il2rgtm1Wjl/SzJ) mice (∼6 weeks old) via tail vein. Three days later, 8 million of the control or IDR CAR T cells in 100 uL PBS were injected by tail vein. In vivo imaging of bioluminescence was performed to monitor tumor growth. Blood was drawn at indicated time points post CAR-T treatment for flow cytometry analysis of CAR-T. To generate solid tumor models, 2.5 million HT29 cells in 100uL PBS were subcutaneously injected into the right flank of NSG mice. Eight days later, when tumor became palpable, 8 million of CAR T cells in 100uL PBS were injected by tail vein, which was followed by a second dose treatment of 4 million of CAR T cells after five days. Tumor growth was measured with calipers every two days, and the size was calculated as one-half of the product of perpendicular length and square width in cubic millimeters (Volume=1/2*L*W*W). Mice were euthanized when the tumor size exceeded 2,000 mm^3^. Blood was collected and tumors were dissected to quantify tumor-infiltrating T cells.

### 8. Signaling characterization in CAR-T

CAR-T cells were resuspend in image medium (RPMI1640 no phenol red supplemented with 20 mM HEPES pH 7.4) and co-cultured with cancer cells at E:T=1:1 for indicated time at 37°C, followed by fixation and permeabilization on ice for 15 min, and staining with individual phosphor-antibodies before sending for flow cytometry analysis or confocal microscopy.

### 9. Microscopy

TIRF and Confocal microscopy was performed on a Nikon Ti2-E inverted motorized microscope stand equipped with motorized stage with stagetop Piezo, Nikon H-TIRF, Yokogawa CSU-X1 spinning disk confocal, Agilent laser combiner with four lines, 405, 488, 561, and 640 nm, and scientific CMOS camera Photometrics Prime 95B. Images were acquired using Nikon Elements. To monitor CAR localization on the cell surface, glass was coated with 10 nM ICAM-1 (Sinobiological #10346-H08H) in PBS for 2 hrs at room temperature. CAR-T cells resuspended in imaging medium (RPMI1640 no phenol red supplemented with 20 mM HEPES pH7.4) were dropped onto the glass and examined by TIRF microscopy. To examine cell-cell conjugates and synapse, CAR-T cells and cancer cells were fixed in imaging media at E:T=1:1 for 30 min at 37°C. The cell mixture was dropped onto glass and imaged by confocal microscopy. To measure signaling in the CAR-T synapse, CAR-T cells were co-cultured with cancer cells for 5 min at 37°C before fixation and permeabilization for 15min on ice, followed by staining with phosphor-antibodies for 30 min on ice. The stained cells were dropped onto glass and imaged by confocal microscopy.

### 10. Image analysis

Microscopy images were analyzed in Fiji (ImageJ). The same brightness and contrast were applied to images within the same panels. CAR condensation was quantified as normalized variance, which equals to the square of standard deviation divided by the mean. Cell conjugation percentage was calculated by dividing the number of tumor cell-associated GFP+ CAR-T cells by the number of total GFP+ CAR T cells. CAR activation was calculated by dividing the intensities of pCD3ζ (Y142) or pLAT (Y171) in the synapse by the intensities of CAR-GFP.

### 11. Statistical analysis

Student’s T test or Mann–Whitney U test analysis was used to assess significance, with P < 0.05 considered significant. Data were analyzed on GraphPad Prism. The statistical details for each experiment were provided in the associated figure legends.

